# Pollinator response diversity and turnover buffer pollination networks from habitat loss in human-modified landscapes

**DOI:** 10.64898/2026.11.20.768076

**Authors:** Nerea Montes-Pérez, Francisco Rodríguez-Sánchez, Ignasi Bartomeus

**Affiliations:** Estación Biológica de Doñana, Department of Ecology and Evolution, Spain; Universidad de Sevilla, Departamento de Biología Vegetal y Ecología, Spain

**Keywords:** Habitat loss, dispersal, spatial connectivity, ecosystem stability, ecological networks, rewiring, pollinators

## Abstract

Agricultural intensification and habitat fragmentation are major drivers of biodiversity loss, yet their effects on ecological interaction networks remain elusive. In plant–pollinator systems, many species disappear with habitat transformation, but network responses remain inconsistent across studies. We hypothesize that this mismatch arises because local observations often overlook the response diversity of the regional species pools. To test this, we compared observed plant–pollinator networks along a land-use gradient with expectations under hypothetical closed-community simulations. Using data from 30 sites in the Doñana Protected Area (southern Spain), we found that spatially connected communities maintained a stable network structure despite habitat loss. This stability was driven by pollinator response diversity, leading to marked species turnover across sites. In contrast, simulated closed communities showed reduced buffering capacity and predominantly negative trends in network metrics such as pollinator richness, total visits, and plant link density. Our results indicate that network stability can emerge from spatial connectivity rather than purely local processes.

## Introduction

Land-use change is a major contemporary threat to biodiversity and ecosystem functioning (Allan *et al*. 2015). Through habitat loss, fragmentation, and landscape simplification, it reduces resource availability and constrains species movement (Fahrig 2003, 2013; Newbold *et al*. 2015). These effects are particularly pronounced for pollinators, which rely on floral resources and suitable nesting habitats (Hellerstein *et al*. 2017; Potts *et al*. 2010). Although some bee species exploit anthropogenic habitats, potentially owing to greater foraging flexibility (Collado *et al*. 2019; Noreika *et al*. 2019), most studies report declines in pollinator abundance and richness with increasing land-use intensification (Didita *et al*. 2025; Millard *et al*. 2021). However, responses are often group-specific (Tsujimoto *et al*. 2022) and show stronger significant trends when extreme habitat loss is included (Winfree *et al*. 2011). Thus, a substantial proportion of the observed variation depends on both the focal taxa and the methodological choices underlying each study.

The observed variability in pollinator community responses to land-use change is partly shaped by landscape context (Martin *et al*. 2019). Habitat heterogeneity can buffer the negative impacts of landscape fragmentation on pollinator richness and abundance by increasing resource diversity across the landscape (Priyadarshana *et al*. 2024). That is, for mobile organisms, the matrix is often permeable, and can be used to different degrees by different species (Ricketts 2001). For example, beyond focal habitat quantity, habitat quality of both the focal and non-focal habitats can further modify pollinator responses to habitat loss (Biegerl *et al*. 2025; Fijen *et al*. 2025). However, species-specific response diversity to changes in land use is rarely quantified (Stavert *et al*. 2017, Cariveau *et al*. 2013)

Changes in species abundances driven by habitat loss can cascade to the interactions they form, reshaping plant–pollinator network structure. Environmental gradients, including land-use intensification, reorganize plant–pollinator networks across landscapes (Proesmans *et al*. 2024; Shinohara *et al*. 2019; Tylianakis & Morris 2017). Habitat loss can modify plant-pollinator network connectance, specialization, and species roles, and may drive interaction loss (Aizen *et al*. 2012; Valiente-Banuet *et al*. 2015) or rewiring (Lázaro & Gómez-Martínez 2022) as natural habitat declines. Moreover, different forms of land-use change and landscape simplification can reduce interaction diversity, potentially weakening network resilience (Gay *et al*. 2024; Maurer *et al*. 2024). However, networks can also be stable, or even increase their robustness against extinctions in crop or urban-dominated landscapes (Proesmans *et al*. 2024; Redhead *et al*. 2018). These structural shifts are critical because network architecture underpins resilience to species loss, pesticide exposure, and other abiotic stressors (Domínguez-García *et al*. 2024; Kaiser-Bunbury *et al*. 2017; Memmott *et al*. 2004). Such contrasting responses suggest that key mechanisms shaping network dynamics may remain unresolved, particularly when the roles of species turnover, dispersal, and regional species pools are not explicitly considered. This is especially challenging because these processes unfold over temporal scales that often exceed the duration of most empirical studies, limiting our ability to detect them directly (Montgomery *et al*. 2020).

To address this limitation, space-for-time substitutions are widely used to infer temporal dynamics from spatial gradients, particularly in fire recovery and community assembly studies (Albrecht *et al*. 2010; Ponisio *et al*. 2017; Potts *et al*. 2010). Although this approach offers practical advantages for insect conservation (Blüthgen *et al*. 2022), it may misrepresent the magnitude or even the direction of change when systems are non-stationary (Damgaard 2019; Perret *et al*. 2024). Importantly, these approaches rarely make explicit how regional species pools contribute to local network dynamics, potentially obscuring mechanisms that buffer ecological functioning. This underscores the need for frameworks integrating spatial connectivity and species-specific habitat preferences shaping interaction assembly (Winfree *et al*. 2014; Collado *et al*. 2019; Gathof *et al*. 2022; Vasiliev & Greenwood 2023). Moving beyond isolated local networks towards spatially connected systems, or meta-networks, provides a conceptual basis to capture these processes by linking local interactions to regional species pools and dispersal dynamics (Chase *et al*. 2020; Hagen *et al*. 2012; Leibold *et al*. 2004). Despite its conceptual promise, this approach has rarely been applied to empirical mutualistic networks, and existing simulations largely focus on general structural properties rather than landscape-driven interaction dynamics (Emer *et al*. 2018; Evans *et al*. 2013; Librán-Embid *et al*. 2021; Rahimi & Jung 2023).

In this study, we assess plant–pollinator network dynamics along a habitat loss gradient, evaluating how pollinators’ response diversity can shape species turnover across sites and drive network stability within an open network framework. Using a comprehensive dataset of plant–pollinator interactions across 30 sites along a land-use gradient, we assess how the surrounding landscape composition shapes pollinator species-specific responses and how this response diversity translates into network reorganization under two contrasting scenarios: with and without species dispersal across habitat patches. To do so, we compare the observed, spatially connected networks with simulated closed-network scenarios with no dispersal allowed among sites. This approach allows us to better capture real-world community dynamics. Specifically, we aim to answer three questions: (i) do wild pollinator responses to landscape composition differ between species?, (ii) does this response diversity lead to stable networks?, and (iii) is this stability maintained by local interaction rewiring or through species turnover from the regional pool? Assuming high dispersal potential and spatial connectivity among habitat patches, we hypothesize that high pollinator response diversity enhances network stability under land use change. Our results demonstrate that integrating a response diversity perspective into a space-for-time design can overcome key limitations of chronosequences, providing a robust framework for future research in the absence of long-term data.

## Methods

### Study area

The study was conducted in the surroundings of the Doñana Protected Area (Southwestern Spain), a heterogeneous Mediterranean landscape characterized by a mosaic of natural scrublands, marshes, and pine forests, embedded in an agricultural matrix. This environmental gradient provides strong variation in the percentage of natural habitat within small spatial scales, making it an ideal system to evaluate how land-use change influences pollinator communities and plant–pollinator interactions.

### Plant–pollinator interaction surveys

We selected sites of Mediterranean shrubland with a similar vegetation composition (i.e., *Salvia rosmarinus*, *Lavandula spp.,* or *Cistus salviifolius*, among others), and spanning a natural-to-agricultural land-use gradient, defined by the proportion of natural habitat within a 2-km radius around each location (Fig. 1(a)). This buffer area was selected taking into account the standard maximum foraging distances previously described in the literature to include a wide range of landscape structure scales (Steffan-Dewenter et al., 2002; Kendall et al., 2022). Plant–pollinator interactions were sampled at each site during four rounds distributed across the flowering season of 2022, capturing early-, mid-, and late-season community activity. Final analyses were done by pooling all rounds, creating a network per site that integrates the seasonal variability.

**Fig. 1.**
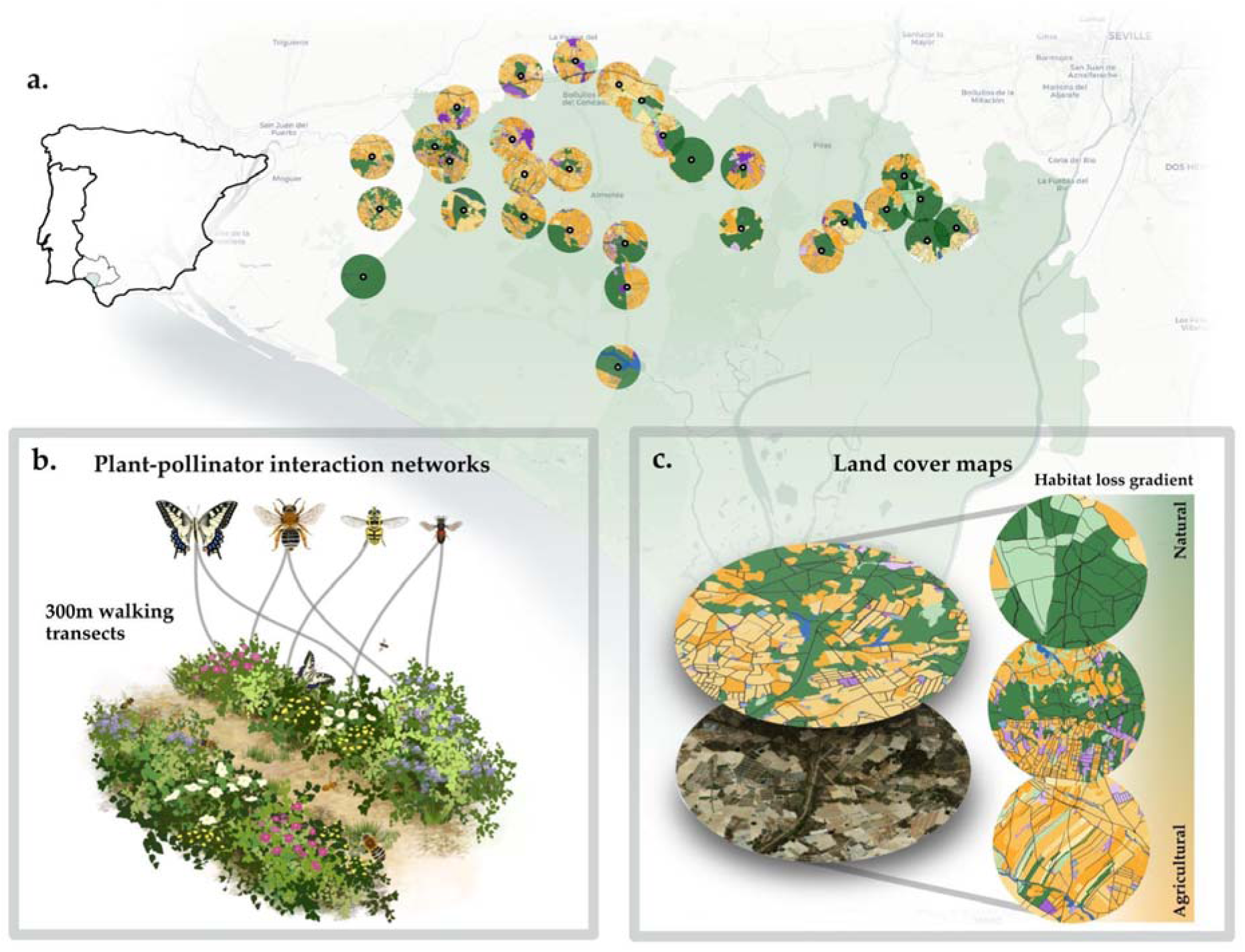
Overview of study design and data collection. a) Site distribution inside and outside of the Doñana Protected Area in green. Sites were placed in seminatural habitat patches with various degrees of habitat loss in the surrounding 2 km buffer. b) Plant pollinator interaction network sampling. c) Creation of land cover maps using satellite imagery. Illustrations and icons were created by an author.

At each site and sampling round, we established standardized 300 m transects and conducted timed flower-visitor observations for 30 effective minutes. Observers walked transects at a constant pace and recorded every legitimate flower visit (i.e., contact with reproductive structures) to an identified plant species. When field identification of insects was not possible, specimens were collected for later identification in the laboratory. Handling time was discounted from the total observation period to ensure that all transect walks had the same effective sampling time. The time of day was randomized across rounds to avoid sampling bias related to pollinator activity patterns. Surveys were conducted only under suitable weather conditions (sunny, no wind, and temperatures above 15 °C).

Simultaneously, we estimated floral resource availability by recording the abundance of plant species in bloom within the transect area. Floral abundance was assessed visually as percentage floral cover for each species, taking into account both its local abundance and its spatial distribution across consecutive 10 m subtransects. Cover estimates therefore represented the overall availability of each flowering species across the transect rather than a value calculated from individual subtransect measurements. Estimated floral cover ranged from 1% to 80%, with higher values corresponding to species that were abundant and/or widely distributed throughout the transect. Rare or highly localized plants were assigned low cover values (1–9%), with 1% representing extremely rare occurrences, such as a single flower across the entire transect.

Lastly, collected insects were pinned or stored appropriately and identified to species or morphospecies by experienced taxonomists (see acknowledgements). All pollinator specimens were resolved to the highest possible taxonomic level; while most pollinator taxa have been identified to the species level, 30% were only identified to the genus level. Hereafter, we will refer to them as pollinator taxa. A list of all pollinator species included can be found in Table S1. Plant species were identified using regional floras. A list of all plant species included can be found in Table S2.

We selected four major, representative pollinator taxa (wild bees, bumblebees, butterflies, and syrphids; excluding Coleoptera and other non-syrphid Diptera) present in at least three sites, so that their response to the land-use gradient could be minimally assessed. We also removed *Apis mellifera,* as its distribution in our region reflects beehive management and not a natural response to landscape features. We only kept plant species that had at least one interaction with any of the selected pollinator taxa. From the original dataset of 270 pollinator species and 86 plant species, we retained 78 pollinator taxa and 56 plant species across 30 sites for the analyses.

### Environmental and land-use variables

Land-use composition surrounding each site was derived from high-resolution land-cover layers created through manual interpretation of satellite imagery accessed via QGIS (QGIS, 2026). We divided land use types into non-managed seminatural areas (Mediterranean shrublands and pine forest), managed seminatural areas (wood plantations, *dehesas,* and road edges), agricultural areas (permanent crops, annual crops, greenhouses, and their associated water reservoirs), and urban areas (buildings, miscellaneous infrastructure, roads, and resource exploitation areas such as mines). Hereafter, natural habitat refers only to non-managed seminatural areas.

For each site, we calculated eight different landscape variables describing multiple axes of landscape complexity and natural habitat loss. The calculated variables are the percentage of seminatural habitat loss (percentage of area without seminatural habitat in the buffer), edge length and density, largest and mean patch size, number of natural habitat patches, habitat diversity, and habitat richness (Table S3). Because the 2 km buffer was selected a priori based on published estimates of maximum pollinator foraging distances, we additionally evaluated the consistency of landscape metrics across spatial extents to assess whether local and broader landscape contexts captured similar patterns. Most metrics showed moderate to strong positive correlations between the 2 km buffer and the 500 m and 1 km buffers (Fig. S1), supporting the use of the 2 km buffer as a descriptor of landscape context in subsequent analyses.

We tested which of these variables was the best predictor of the aggregated pollinator species visitation rates per site by comparing Akaike’s Information Criterion (AIC) values. The percentage of seminatural habitat loss was the best predictor and was thus used in all subsequent analyses as the main predictor of land-use intensity (Table S4).

### Statistical analyses

Our statistical analyses included three major steps. In summary, we first analysed observed changes in network properties and community composition across the 30 sites. Second, we assessed taxon-specific pollinator responses to the percentage of natural habitat loss in a 2km radius buffer to characterize the response of each pollinator to land-use change. Lastly, we simulated a closed-community scenario in which pollinator immigration was removed, thus constraining species turnover and only allowing interaction rewiring to occur. In this scenario, networks were constrained to the original pool of pollinator taxa and plant species for each site, and predictions of interaction abundance along a habitat loss gradient were done only on those original species, preventing both the reintroduction of extinct species and the arrival of new ones. This simulation design allows us to evaluate how network dynamics would change driven only by local interaction rewiring in the absence of pollinator dispersal (see below for the reconstruction of species interactions in the simulation).

### Dynamics of open communities and the metanetwork

We first analysed observed network properties to evaluate the resilience of open communities to habitat loss. For each empirical network, we calculated pollinator richness, pollinator link density, plant link density, connectance, nestedness (NODFc: NODF corrected for matrix size; Song *et al*. 2017), and the total number of visits. We selected these metrics because they are easily interpretable and have clear theoretical links to network robustness, interaction reorganization, and resilience under species loss and environmental disturbance (Aizen *et al*. 2012; Memmott *et al*. 2004; Poisot *et al*. 2012b). We fitted generalized additive models (GAMs; Wood 2017) to model the variation of each metric along the habitat-loss gradient, using the metric as the response variable and a smooth term (thin plate regression spline) of habitat-loss percentage as the predictor. To evaluate statistically significant changes along the curves, we also performed posterior simulations of the first derivatives to identify sections of the gradient where the slope differed significantly from zero (package *gratia*; Simpson 2024). This approach allowed us to compare the dynamics of the observed open-community networks with those obtained under the simulated closed scenario described in the section below.

We next calculated the two main axes of change that could drive network stability and interaction reorganization: species turnover and interaction dissimilarity. We first quantified pollinator compositional shifts (number of observed visits) using Bray–Curtis dissimilarity using the function “vegdist” from *vegan* package (Oksanen *et al*. 2025). Then, we assessed interaction dissimilarity driven by interaction rewiring (Poisot *et al*. 2012b) using the “betalinkr” function from *bipartite* package (Dormann *et al*. 2009). For interaction dissimilarity, we focus exclusively on interaction rewiring among shared species, as changes due to species turnover are better assessed by looking at compositional turnover.

### Pollinator visitation responses to landscape variables

Secondly, we assessed taxon-specific pollinator visitation responses to landscape configuration. We used generalized additive mixed models (GAMMs; Pedersen et al. 2019) to evaluate non-linear responses of pollinator visits to landscape composition, summing all visits performed by each pollinator within each observed network. Following Mainguy et al. 2026 and the recommendations of the *mgcv* package (Wood 2011, 2017), we first fitted candidate models using maximum likelihood (ML) to enable model comparison and selection of optimal landscape variables and buffer size based on Akaike’s Information Criterion (AIC). The best-supported model was then refitted using restricted maximum likelihood (REML) to obtain more reliable smoothness estimates and reduce bias. Further details on model selection are provided in Table S4. Pollinator visitation was modelled following a zero-inflated Poisson distribution and a taxon-specific smooth function of landscape composition:

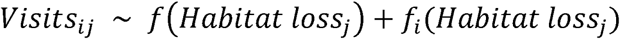

Where *Visits_ij_* represents the number of visits of pollinator *i* at site *j*,*f*(*Habitat loss_j_*) describes the average community-level response to habitat loss, and *f_i_*(*Habitat loss_j_*) represents pollinator-specific deviations from this general trend. This formulation allows individual pollinator taxa to exhibit distinct non-linear responses to habitat loss while simultaneously estimating the average response across the pollinator community (Pedersen et al. 2019). For plant species, the same modelling approach was applied to flower abundance, but no consistent response to landscape variables was detected. This is expected, given that the study design targeted focal habitats with different landscape contexts, but similar plant composition.

Additionally, we evaluated pollinator visitation curves to quantify net changes along the habitat-loss gradient and to estimate the number of species showing positive or negative trends at each 1% habitat-loss step, in order to detect potential critical shifts in the community. To do this, we calculated the first derivative of each pollinator visitation curve at every 1% step. Finally, for each pollinator taxon, we calculated a net trend by summing the first-derivative estimates across the land-use gradient. The direction of this value, represented by colour, is provided only as a visual reference of the taxon’s potential overall trend. It should not be interpreted as a definitive assessment, as responses to land-use change are nonlinear and reducing them to a single value may obscure important variation along the gradient. Additionally, we summarised net trends per pollinator family to obtain a general overview of habitat loss impact across major pollinator groups (Supplementary Figure S2).

### Closed-communities simulations

Thirdly, we simulated a closed-community scenario in which the species composition at each site was restricted to that of the original observed networks, and pollinator visits were predicted along the habitat-loss gradient according to the taxon-specific GAMM model. In particular, we used the results of the visitation curves for each pollinator taxa and the predicted total number of visits performed by each pollinator at 5% steps of habitat loss. The predicted number of visits of each pollinator per habitat loss step was distributed among the plant species based on the empirically observed visitation proportions of each pollinator to each plant species, allowing us to reconstruct interaction networks with predicted visitation frequencies. Plant species composition remained constant within each observed network. When a pollinator species dropped below 0.5 visits per habitat loss step, we considered it extinct. This threshold was used to avoid premature extinctions, as predicted visitation values were slightly underestimated by the GAM. Once a pollinator species became extinct, it was not allowed to reappear in later steps, as this would represent reintroduction through migration, which is not permitted in the closed-community scenario. For each simulated network, we calculated the same set of network metrics as in the open-community scenario, allowing direct comparison of network dynamics between the two approaches.

We performed all analyses using R software version 4.5.1 (R Core Team, 2025). Model checks were done with the DHARMa package (Hartig 2024). We assessed spatial autocorrelation in observed network metrics, pollinator diversity, and the residuals of the pollinator habitat preference models using Moran I index (package spdep; Bivand et al. 2011). Network metrics and model residuals did not exhibit significant spatial autocorrelation.

## Results

### Dynamics of observed communities and their networks

The pollination interaction networks obtained for each of the 30 sites showed no consistent impact of natural habitat loss on their structure and properties (Fig. 2). The only metric showing statistically significant trends along the gradient is nestedness, which increases along the gradient (Fig. 2d). On the other hand, trends for all other metrics, such as connectance or total visits, although slightly positive or negative, were not statistically significant.

**Fig. 2.**
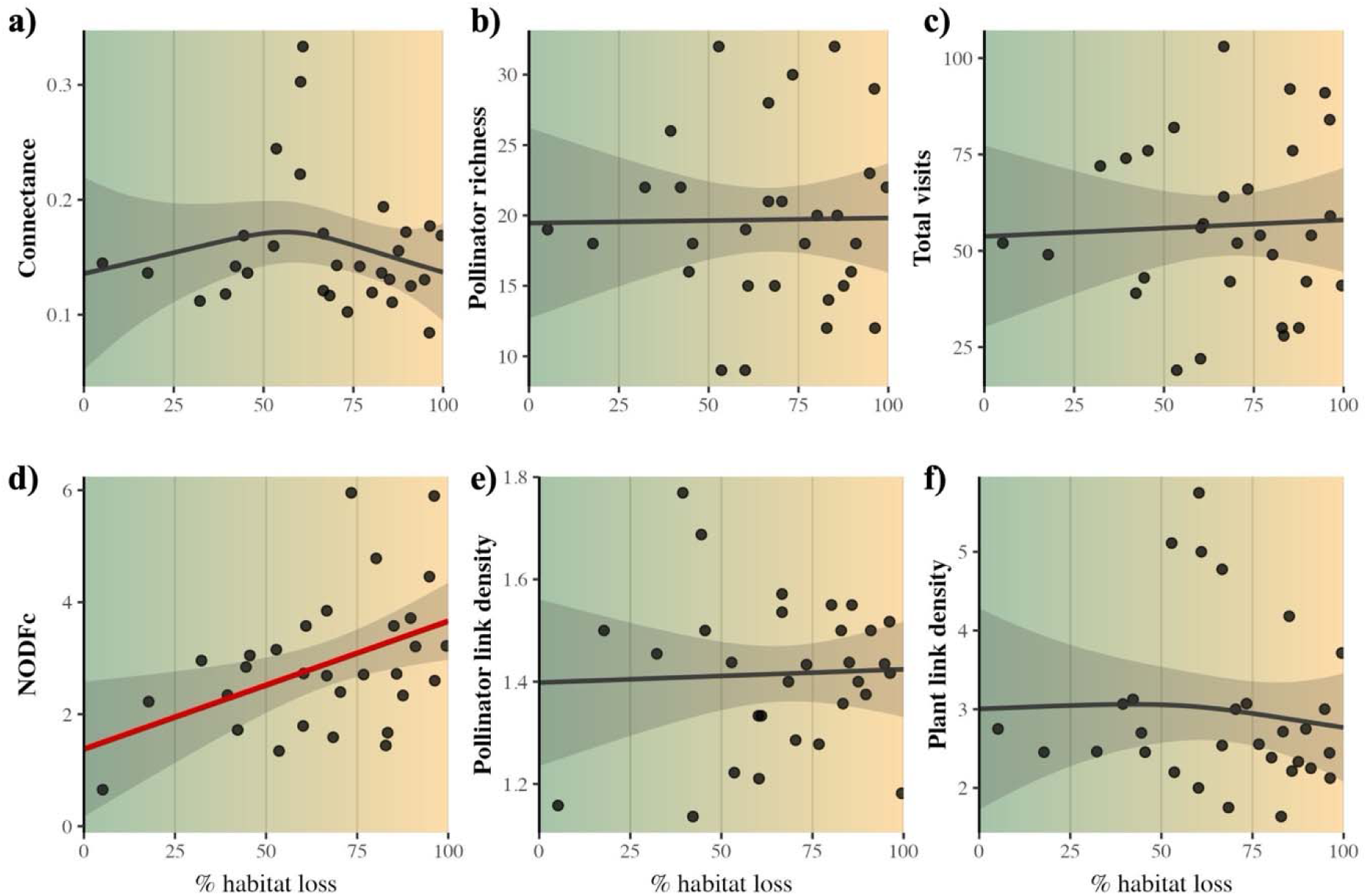
Observed plant–pollinator network metrics along the habitat-loss gradient. Panels, read left to right and then top to bottom: (a) connectance, (b) pollinator richness, (c) total visits, (d) NODFc (nestedness corrected for matrix size), (e) pollinator link density and (f) plant link density. The green-to-orange background represents increasing habitat loss within a 2 km radius. Lines show fitted generalised additive model trends with shaded uncertainty; red indicates a significant trend.

Overall, pollinator composition showed a high turnover of taxa and low rewiring between shared species, with no significant pattern along the gradient. In particular, the Bray–Curtis beta-diversity index suggested a gradual decrease in species turnover between pairs of adjacent sites ordered along the gradient (i.e. site 1 vs site 2, site 2 vs site 3, and so forth; Fig. 3 (a)). Interaction rewiring between shared species along the gradient is overall low and barely increases with habitat loss (Fig. 3 (b)).

**Fig. 3.**
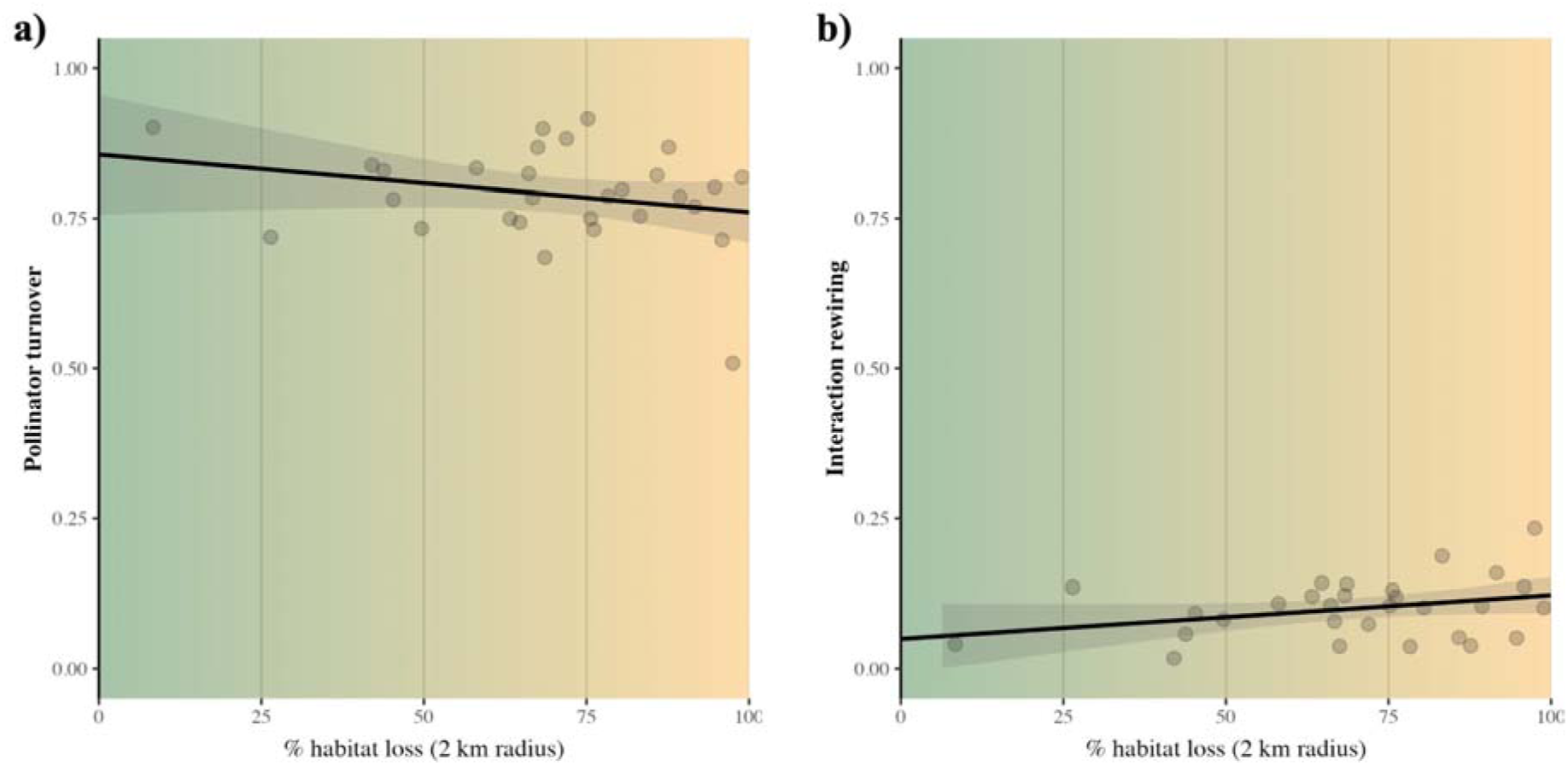
Community composition changes and interaction dissimilarity along the habitat loss gradient between consecutive sites. (a) Bray Curtis betadiversity index and (b) interaction rewiring among shared species (Beta OS). Background colours from green to orange represent 100% natural landscapes up to 100% agricultural landscapes. Black lines show statistically non-significant trends.

### Pollinator visitation responses to habitat loss

Responses to habitat loss varied substantially in shape, magnitude and direction among the 78 pollinator taxa analysed (Fig. 4a), with some taxa declining and others showing non-linear or increasing trends along the gradient. For instance, flies such as *Usia sp*. and *Rhyncomia cuprea* showed consistent increases in predicted visits with habitat loss, indicating an association with more disturbed agricultural landscapes. In contrast, bees such as *Nomada succincta* and *Andrena morio* showed consistent decreases, with more visits predicted in landscapes containing a higher proportion of natural habitat (Fig. S2). These contrasting patterns highlight the diversity of responses both among taxa and across pollinator guilds. Overall, however, most pollinator taxa did not show strong declines in disturbed habitats.

**Fig. 4.**
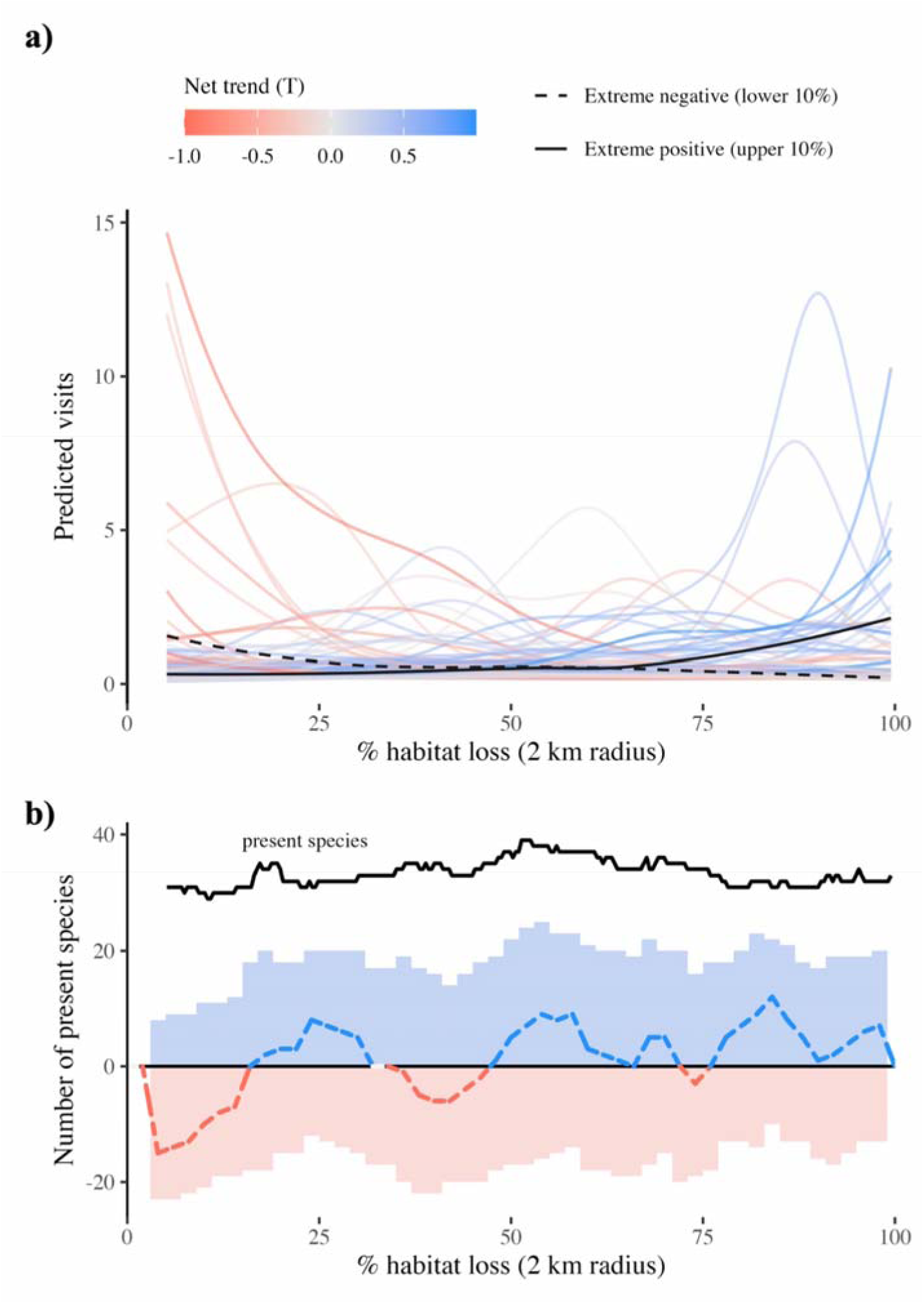
Pollinator responses to the habitat-loss gradient. (a) Predicted visits for 78 pollinator taxa from generalised additive mixed models, displayed on a pseudo-logarithmic scale. Curve colour represents each taxon’s net trend across the gradient (red, negative; blue, positive); a dashed black curve highlights the lower 10% and a solid black curve the upper 10% of net trends. These curves may cross because individual responses are nonlinear, while the ranking reflects the entire gradient. (b) The black line shows the predicted number of present taxa (at least 0.5 visits); blue and red shading show counts of taxa with positive and negative local slopes, respectively, at each 1% habitat-loss step. The coloured dashed line is their net balance (positive minus negative taxa).

We also identified critical points along the gradient characterized by increased losses of pollinator taxa (Fig. 4 (b)), particularly in highly natural habitats and in intermediate agricultural landscapes with half of the natural habitat available (40 to 65% habitat loss). Pollinator species richness increased moderately with habitat loss (Spearman’s ρ = 0.19 at the 2 km buffer).

### Closed communities scenarios

When simulating the effect of increasing habitat loss but no pollinator immigration for each site (i.e. closed communities), we found that habitat loss negatively impacted many network structural indices and community properties (Fig. 5). As expected, pollinator richness, total visits, and plant link density showed a strong negative impact as habitat loss increases. In contrast, connectance, nestedness and pollinator link density showed much weaker trends, except for the most transformed habitats (between 75% and 100% habitat loss).

**Fig. 5.**
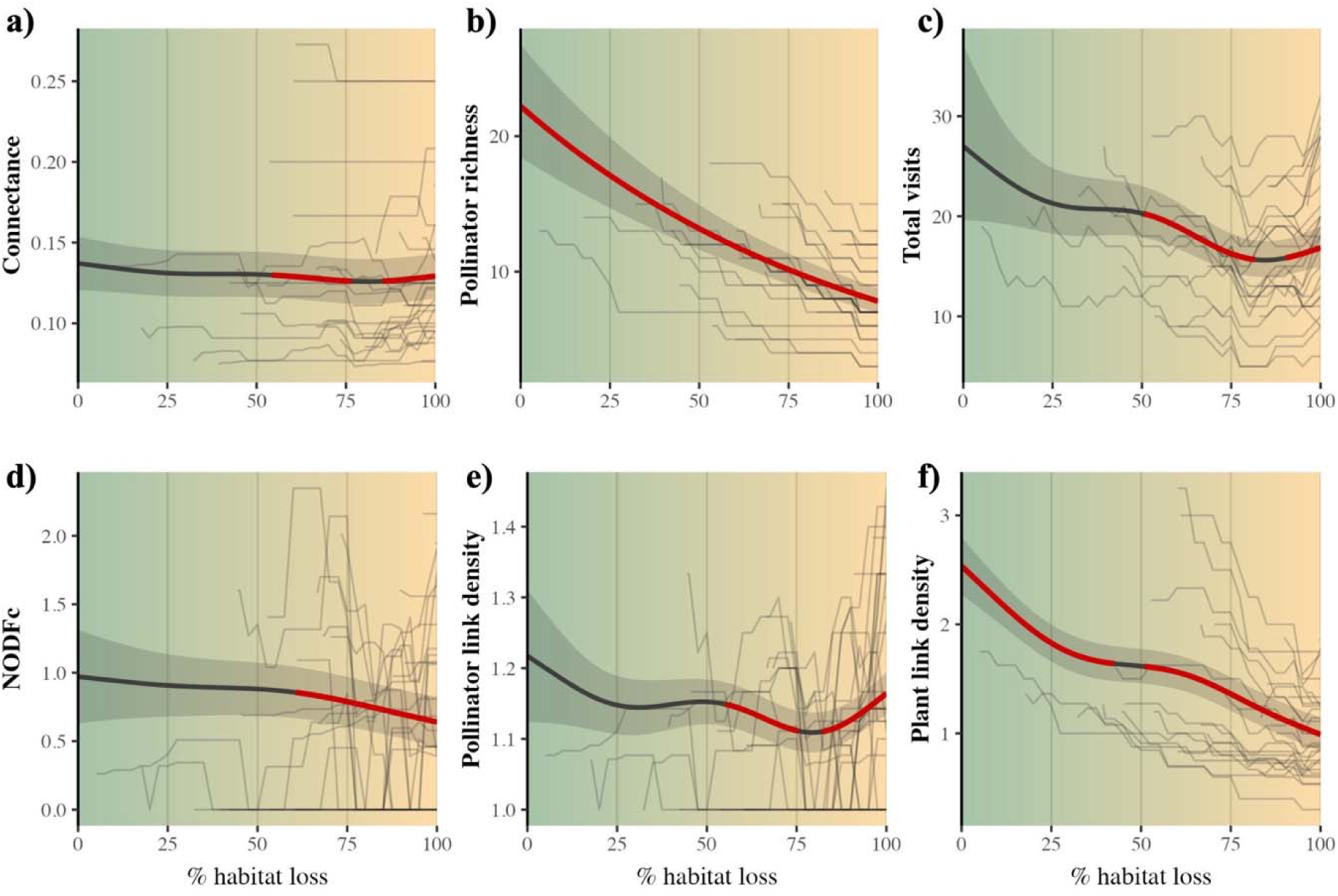
Simulated closed-community network metrics under increasing habitat loss, with no pollinator immigration. Panels, read left to right and then top to bottom: connectance, pollinator richness, total visits, NODFc, pollinator link density and plant link density. Thin lines show site-specific trajectories and thick lines show the average trajectory across sites; red segments indicate slopes significantly different from zero.

## Discussion

Our results show that plant–pollinator networks in open, real-world communities are primarily shaped by pollinator response diversity and their ability to disperse across the landscape. The marked variation in responses to habitat loss across pollinator taxa may buffer pollination networks against collapse. This suggests that partial landscape disturbance may promote community reassembly rather than generalized pollinator decline, potentially favoring disturbance-adapted species (Stavert *et al*. 2017). This reshuffling of species composition emerges as the dominant mechanism underlying network dynamics along the gradient, buffering the negative effects typically associated with habitat loss. In contrast, when communities are simulated as closed systems without dispersal or species replacement, this buffering effect disappears and network structure deteriorates rapidly. Together, these findings highlight species turnover and dispersal under a metanetwork framework as key processes maintaining network stability impacted by landscape transformation.

Pollinator composition showed consistently high dissimilarity across the habitat loss gradient, indicating substantial species turnover among consecutive communities across the land use gradient. Although dissimilarity tended to decrease slightly towards more degraded sites, this pattern was not statistically significant, suggesting that high compositional turnover was maintained across the gradient. Despite this turnover, most network-level properties remained remarkably stable for the observed communities. These findings align with studies focused on temporal reorganizations showing that ecological networks can remain structurally stable over time despite substantial interannual turnover in species composition (CaraDonna et al. 2017; Chacoff et al. 2012; Domínguez-Garcia et al. 2026). This decoupling between species turnover and network structure suggests that networks can maintain their overall architecture even as species identities change (Kaiser-Bunbury et al. 2017; Memmott et al. 2004). Such stability contrasts with the expectation that habitat loss leads to systematic degradation of ecological networks through species loss and interaction breakdown (Aizen et al. 2012; Valiente-Banuet et al. 2015), and instead supports the view that network properties can be relatively robust to compositional change when species responses are heterogeneous (Proesmans et al. 2024; Winfree et al. 2014). Together, these spatial and temporal perspectives suggest that turnover-driven stability may be a general property of ecological networks. But maintaining such turnover requires a high diversity of species responses, allowing pollinators to replace one another across space and time.

Pollinators exhibited clear taxon-specific responses to land-use change. While some taxa declined (mainly bees and syrphids), many persisted or even increased in disturbed habitats (e.g. some butterflies and other fly families), reflecting differences in ecological traits, resource use, and tolerance to environmental change (Stavert et al. 2017; Noreika et al. 2019; Tsujimoto et al. 2022). These taxon-specific responses are reflected in high pollinator turnover but stable pollinator richness along the habitat loss gradient. Previous studies show mixed evidence about the impacts of habitat fragmentation on pollinator richness and abundance. On one hand, many studies have reported a decrease in pollinator richness and abundance with natural habitat loss (Olhnuud et al. 2025), at least when habitat loss is extreme (Winfree et al. 2009). On the other hand, increasing habitat heterogeneity as natural habitat declines may promote higher species richness (Parreño et al. 2024; Priyadarshana et al. 2024), and, consequently, greater species turnover. This indicates that anthropogenized landscapes can act not only as filters but also as arenas for community reassembly, where species are redistributed rather than uniformly lost.

Importantly, interaction rewiring among shared species was present but limited and did not show any consistent pattern along the gradient. This suggests that network reorganization is driven primarily by species replacement rather than behavioural flexibility in the interactions of co-occurring species (Domínguez-Garcia et al. 2026). Such a pattern is expected, as high levels of species turnover inevitably lead to high interaction turnover (Poisot et al. 2012a). Although rewiring has been proposed as a key mechanism of network resilience (CaraDonna et al. 2017), particularly across spatial rather than temporal gradients (Magrach et al. 2023), and may increase under habitat loss (Lázaro & Gómez-Martínez 2022), many of these studies focus on within-season dynamics or do not explicitly account for landscape variables. In contrast, our results suggest that when processes underpinning network stability depend strongly on landscape structure, with a large species turnover, the role of interaction rewiring is secondary. This is particularly relevant given the importance of landscape connectivity for pollinator dispersal (Vasiliev & Greenwood 2023). However, note that a low degree of rewiring among the few shared species in our networks does not diminish the importance of behaviour flexibility, as the generalized nature of plant-pollinator interactions ensures the integration of new species in the network (Waser et al. 1996).

Within a metanetwork framework, local communities can be viewed as subsets of a regional pool of interacting species, continuously reassembled through dispersal (Chase et al. 2020; Hagen et al. 2012; Leibold et al. 2004). Although we know very little about pollinator dispersal rates, the few pieces of evidence indicate that even for taxa that are central nest foragers and have strong philopatry, dispersal of several km per year is not infrequent (Dorian & Crone 2026). The combination of dispersal and high response diversity among pollinators allows functionally similar species to substitute for one another across sites, maintaining key network properties (Blüthgen et al. 2008; Poisot et al. 2012a; Stavert et al. 2017). Although empirical work explicitly linking landscape structure to network dynamics remains limited, existing studies indicate that landscape composition and connectivity can shape both species distributions and interaction patterns (Pellissier et al. 2018; Trøjelsgaard & Olesen 2016; Tylianakis et al. 2010). This interpretation is supported by our simulated closed communities, where preventing dispersal and species replacement led to strong declines in richness, interaction frequency, and network structure. Additionally, connectance and pollinator link density remained largely unchanged along the gradient. This stability may have emerged because the reduction in interactions was proportional to the loss of pollinator species, despite a slight increase in both metrics at the section of the gradient with the strongest species turnover. Because land-use gradients are embedded within broader landscape contexts, isolating the effects of habitat loss from co-varying environmental factors remains challenging. Nevertheless, our results demonstrate that the apparent robustness of plant– pollinator networks depends critically on their openness and connectivity to surrounding communities.

## Conclusion

Overall, our findings suggest that maintaining landscape connectivity and access to regional species pools is likely essential for sustaining plant–pollinator interactions under ongoing land-use change. While local habitat degradation reshapes communities, dispersal-mediated turnover can buffer network structure and maintain ecosystem functioning. These dispersal processes are mediated by pollinator responses to habitat loss, which allow for a continuous species turnover as habitat is lost. Such stability of network properties is removed when we impede pollinator dispersal under a closed community scenario. Hence, this buffering capacity may depend on the persistence of functionally diverse species and sufficiently connected landscapes, and may break down under more extreme or rapid habitat loss or fragmentation. These results highlight the importance of integrating spatial processes and meta-network dynamics into conservation strategies aimed at preserving ecological interactions.

## Data accessibility

The data and code supporting the findings of this study can be found in the supplementary materials and will be publicly available through Zenodo upon acceptance.

## References

Aizen, M., Sabatino, M. & Tylianakis, J. (2012). Specialization and Rarity Predict Nonrandom Loss of Interactions from Mutualist Networks. Science, 335, 1486–9.

Albrecht, M., Riesen, M. & Schmid, B. (2010). Plant–pollinator network assembly along the chronosequence of a glacier foreland. Oikos, 119, 1610–1624.

Allan, E., Manning, P., Alt, F., Binkenstein, J., Blaser, S., Blüthgen, N., et al. (2015). Land use intensification alters ecosystem multifunctionality via loss of biodiversity and changes to functional composition. Ecol. Lett., 18, 834–843.

Biegerl, C., Holzschuh, A., Tanner, B., Sponsler, D., Krauss, J., Zhang, J., et al. (2025). Landscape management can foster pollinator richness in fragmented high-value habitats. Proc. R. Soc. B Biol. Sci., 292, 20242686.

Bivand, R., Anselin, L., Berke, O., Bernat, A., Carvalho, M., Chun, Y., et al. (2011). spdep: Spatial dependence: weighting schemes, statistics and models. R Package Version 05–31 URL HttpCRAN R-Proj. Orgpackage Spdep.

Blüthgen, N., Fründ, J., Vázquez, D.P. & Menzel, F. (2008). What Do Interaction Network Metrics l Us About Specialization and Biological Traits. Ecology, 89, 3387–3399.

Blüthgen, N., Staab, M., Achury, R. & Weisser, W.W. (2022). Unravelling insect declines: can space replace time? Biol. Lett., 18, 20210666.

CaraDonna, P.J., Petry, W.K., Brennan, R.M., Cunningham, J.L., Bronstein, J.L., Waser, N.M., et al. (2017). Interaction rewiring and the rapid turnover of plant–pollinator networks. Ecol. Lett., 20, 385–394.

Cariveau, D. P., Williams, N. M., Benjamin, F. E., & Winfree, R. (2013). Response diversity to land use occurs but does not consistently stabilise ecosystem services provided by native pollinators. Ecology letters, 16(7), 903–911.

Chacoff, N.P., Vázquez, D.P., Lomáscolo, S.B., Stevani, E.L., Dorado, J. & Padrón, B. (2012). Evaluating sampling completeness in a desert plant–pollinator network. J. Anim. Ecol., 81, 190–200.

Chase, J.M., Jeliazkov, A., Ladouceur, E. & Viana, D.S. (2020). Biodiversity conservation through the lens of metacommunity ecology. Ann. N. Y. Acad. Sci., 1469, 86–104.

Collado, M.Á., Sol, D. & Bartomeus, I. (2019). Bees use anthropogenic habitats despite strong natural habitat preferences. Divers. Distrib., 25, 924–935.

Damgaard, C. (2019). A Critique of the Space-for-Time Substitution Practice in Community Ecology. Trends Ecol. Evol., 34, 416–421.

Didita, M., Nemomissa, S., Lemessa, D. & Bekele, T. (2025). The diversity of pollinators within different land use types across agricultural landscapes is influenced by the proximity of surrounding forests. Glob. Ecol. Conserv., 62, e03776.

Domínguez-Garcia, V., Molina, F.P., Allen-Perkins, A., Godoy, O. & Bartomeus, I. (2026). Plant-Pollinator Interaction Rewiring Boosts Year-to-Year Community Persistence. Ecol. Lett., 29, e70293.

Domínguez-Garcia, V., Molina, F.P., Godoy, O. & Bartomeus, I. (2024). Interaction network structure explains species’ temporal persistence in empirical plant–pollinator communities. Nat. Ecol. Evol., 8, 423–429.

Dorian, N. N., & Crone, E. E. (2026). Fat tailed dispersal shapes connectivity in a solitary bee. Ecology, 107(7), e70437.

Dormann, C.F., Fründ, J., Blüthgen, N. & Gruber, B. (2009). Indices, graphs and null models: analyzing bipartite ecological networks.

Emer, C., Galetti, M., Pizo, M.A., Guimarães Jr., P.R., Moraes, S., Piratelli, A., et al. (2018). Seed-dispersal interactions in fragmented landscapes – a metanetwork approach. Ecol. Lett., 21, 484–493.

Evans, D.M., Pocock, M.J.O. & Memmott, J. (2013). The robustness of a network of ecological networks to habitat loss. Ecol. Lett., 16, 844–852.

Fahrig, L. (2003). Effects of Habitat Fragmentation on Biodiversity. Annu. Rev. Ecol. Evol. Syst., 34, 487–515.

Fahrig, L. (2013). Rethinking patch size and isolation effects: the habitat amount hypothesis. J. Biogeogr., 40, 1649–1663.

Fijen, T.P.M., Bishop, G.A., Ganuza, C., Scheper, J. & Kleijn, D. (2025). Analyzing the relative importance of habitat quantity and quality for boosting pollinator populations in agricultural landscapes. Conserv. Biol., 39, e14317.

Gathof, A.K., Grossmann, A.J., Herrmann, J. & Buchholz, S. (2022). Who can pass the urban filter? A multi-taxon approach to disentangle pollinator trait–environmental relationships. Oecologia, 199, 165–179.

Gay, C., Gaba, S. & Bretagnolle, V. (2024). The structure of plant–pollinator networks is affected by crop type in a highly intensive agricultural landscape. Agric. Ecosyst. Environ., 359, 108759.

Hagen, M., Kissling, W.D., Rasmussen, C., De Aguiar, M.A.M., Brown, L.E., Carstensen, D.W., et al. (2012). Biodiversity, species interactions and ecological networks in a fragmented world. In: Advances in ecological research (eds. Dumbrell, A. & Milner, A.). Academic Press, Cambridge, MA (USA), pp. 89–210.

Hartig, F. (2024). DHARMa: Residual Diagnostics for Hierarchical (Multi-Level / Mixed) Regression Models.

Hellerstein, D., Hitaj, C., Smith, D. & Davis, A. (2017). Land Use, Land Cover, and Pollinator Health: A Review and Trend Analysis.

Kaiser-Bunbury, C.N., Mougal, J., Whittington, A.E., Valentin, T., Gabriel, R., Olesen, J.M., et al. (2017). Ecosystem restoration strengthens pollination network resilience and function. Nature, 542, 223–227.

Lázaro, A. & Gómez-Martínez, C. (2022). Habitat loss increases seasonal interaction rewiring in plant–pollinator networks. Funct. Ecol., 36, 2673–2684.

Leibold, M.A., Holyoak, M., Mouquet, N., Amarasekare, P., Chase, J.M., Hoopes, M.F., et al. (2004). The metacommunity concept: a framework for multi-scale community ecology. Ecol. Lett., 7, 601–613.

Librán-Embid, F., Grass, I., Emer, C., Ganuza, C. & Tscharntke, T. (2021). A plant–pollinator metanetwork along a habitat fragmentation gradient. Ecol. Lett., 24, 2700–2712.

Magrach, A., Artamendi, M., Lapido, P.D., Parejo, C. & Rubio, E. (2023). Indirect interactions between pollinators drive interaction rewiring through space. Ecosphere, 14, e4521.

Mainguy, J., McInerney, R., Millar, R.B., Valiquette, E., Bélanger, M. & Moral, R. de A. (2026). Generalized Additive Modeling of Ecological Data With mgcv: New Adequacy Assessment Tools. Ecol. Evol., 16, e72825.

Martin, E.A., Dainese, M., Clough, Y., Báldi, A., Bommarco, R., Gagic, V., et al. (2019). The interplay of landscape composition and configuration: new pathways to manage functional biodiversity and agroecosystem services across Europe. Ecol. Lett., 22, 1083–1094.

Maurer, C., Martínez-Núñez, C., Dominik, C., Heuschele, J., Liu, Y., Neumann, P., et al. (2024). Landscape simplification leads to loss of plant–pollinator interaction diversity and flower visitation frequency despite buffering by abundant generalist pollinators. Divers. Distrib., 30, e13853.

Memmott, J., Waser, N.M. & Price, M.V. (2004). Tolerance of pollination networks to species extinctions. Proc. R. Soc. Lond. B Biol. Sci., 271, 2605–2611.

Millard, J., Outhwaite, C.L., Kinnersley, R., Freeman, R., Gregory, R.D., Adedoja, O., et al. (2021). Global effects of land-use intensity on local pollinator biodiversity. Nat. Commun., 12, 2902.

Montgomery, G.A., Dunn, R.R., Fox, R., Jongejans, E., Leather, S.R., Saunders, M.E., et al. (2020). Is the insect apocalypse upon us? How to find out. Biol. Conserv., 241, 108327.

Newbold, T., Hudson, L.N., Hill, S.L.L., Contu, S., Lysenko, I., Senior, R.A., et al. (2015). Global effects of land use on local terrestrial biodiversity. Nature, 520, 45–50.

Noreika, N., Bartomeus, I., Winsa, M., Bommarco, R. & Öckinger, E. (2019). Pollinator foraging flexibility mediates rapid plant-pollinator network restoration in semi-natural grasslands. Sci. Rep., 9, 15473.

Oksanen, J., Simpson, G.L., Blanchet, F.G., Kindt, R., Legendre, P., Minchin, P.R., et al. (2025). Vegan: community ecology package.

Olhnuud, A., Wen, J., Yu, J., Lyu, F. & Zhang, Q. (2025). Responses of insect pollinators to habitat fragmentation: A global meta-analysis. J. Appl. Ecol., 62, 2502–2514.

Page, M.L. & Williams, N.M. (2023). Evidence of exploitative competition between honey bees and native bees in two California landscapes. J. Anim. Ecol., 92, 1802–1814.

Parreño, M.A., Werle, S., Buydens, L., Leroy, C., Roberts, S., Koirala, S., et al. (2024). Landscape heterogeneity correlates with bee and pollen diversity while size and specialization degree explain species-specific responses of wild bees to the environment. Sci. Total Environ., 954, 176595.

Pellissier, L., Albouy, C., Bascompte, J., Farwig, N., Graham, C., Loreau, M., et al. (2018). Comparing species interaction networks along environmental gradients. Biol. Rev., 93, 785–800.

Perret, D.L., Evans, M.E.K. & Sax, D.F. (2024). A species’ response to spatial climatic variation does not predict its response to climate change. Proc. Natl. Acad. Sci., 121, e2304404120.

Poisot, T., Canard, E., Mouillot, D., Mouquet, N. & Gravel, D. (2012a). The dissimilarity of species interaction networks. Ecol. Lett., 15, 1353–1361.

Poisot, T., Canard, E., Mouquet, N. & Hochberg, M.E. (2012b). A comparative study of ecological specialization estimators. Methods Ecol. Evol., 3, 537–544.

Ponisio, L.C., Gaiarsa, M.P. & Kremen, C. (2017). Opportunistic attachment assembles plant– pollinator networks. Ecol. Lett., 20, 1261–1272.

Potts, S.G., Biesmeijer, J.C., Kremen, C., Neumann, P., Schweiger, O. & Kunin, W.E. (2010). Global pollinator declines: trends, impacts and drivers. Trends Ecol. Evol., 25, 345–353.

Priyadarshana, T.S., Martin, E.A., Sirami, C., Woodcock, B.A., Goodale, E., Martínez-Núñez, C., et al. (2024). Crop and landscape heterogeneity increase biodiversity in agricultural landscapes: A global review and meta-analysis. Ecol. Lett., 27, e14412.

Proesmans, W., Felten, E., Laurent, E., Albrecht, M., Cyrille, N., Labonté, A., et al. (2024). Urbanisation and agricultural intensification modulate plant–pollinator network structure and robustness. Funct. Ecol., 38, 628–641.

Rahimi, E. & Jung, C. (2023). Plant–pollinator metanetworks in fragmented landscapes: a simulation study. Ecol. Process., 12, 29.

Redhead, J.W., Woodcock, B.A., Pocock, M.J.O., Pywell, R.F., Vanbergen, A.J. & Oliver, T.H. (2018). Potential landscape-scale pollinator networks across Great Britain: structure, stability and influence of agricultural land cover. Ecol. Lett., 21, 1821–1832.

Ricketts, T.H. (2001). The Matrix Matters: Effective Isolation in Fragmented Landscapes. Am. Nat., 158, 87–99.

Shinohara, N., Uchida, K. & Yoshida, T. (2019). Contrasting effects of land-use changes on herbivory and pollination networks. Ecol. Evol., 9, 13585–13595.

Simpson, G.L. (2024). gratia: An R package for exploring generalized additive models. ArXiv Prepr. ArXiv240619082.

Song, C., Rohr, R.P. & Saavedra, S. (2017). Why are some plant–pollinator networks more nested than others? J. Anim. Ecol., 86, 1417–1424.

Stavert, J.R., Pattemore, D.E., Gaskett, A.C., Beggs, J.R. & Bartomeus, I. (2017). Exotic species enhance response diversity to land-use change but modify functional composition. Proc. R. Soc. B Biol. Sci., 284, 20170788.

Trøjelsgaard, K. & Olesen, J.M. (2016). Ecological networks in motion: micro- and macroscopic variability across scales. Funct. Ecol., 30, 1926–1935.

Tsujimoto, S.G., Kim, J.Y., Noda, A., Hiratsuka, Y.L. & Nishihiro, J. (2022). Landscape effects on pollinator abundance differ among taxonomic groups. Ecol. Res., n/a.

Tylianakis, J.M., Laliberté, E., Nielsen, A. & Bascompte, J. (2010). Conservation of species interaction networks. Biol. Conserv., Conserving complexity: Global change and community-scale interactions, 143, 2270–2279.

Tylianakis, J.M. & Morris, R.J. (2017). Ecological Networks Across Environmental Gradients. Annu. Rev. Ecol. Evol. Syst., 48, 25–48.

Valiente-Banuet, A., Aizen, M.A., Alcántara, J.M., Arroyo, J., Cocucci, A., Galetti, M., et al. (2015). Beyond species loss: the extinction of ecological interactions in a changing world. Funct. Ecol., 29, 299–307.

Vasiliev, D. & Greenwood, S. (2023). The role of landscape connectivity in maintaining pollinator biodiversity needs reconsideration. Biodivers. Conserv., 32, 3765–3790.

Waser, N. M., Chittka, L., Price, M. V., Williams, N. M., & Ollerton, J. (1996). Generalization in pollination systems, and why it matters. Ecology, 77(4), 1043–1060.

Winfree, R., Aguilar, R., Vázquez, D.P., LeBuhn, G. & Aizen, M.A. (2009). A meta-analysis of bees’ responses to anthropogenic disturbance. Ecology, 90, 2068–2076.

Winfree, R., Bartomeus, I. & Cariveau, D.P. (2011). Native Pollinators in Anthropogenic Habitats. Annu. Rev. Ecol. Evol. Syst., 42, 1–22.

Winfree, R., Williams, N.M., Dushoff, J. & Kremen, C. (2014). Species Abundance, Not Diet Breadth, Drives the Persistence of the Most Linked Pollinators as Plant-Pollinator Networks Disassemble. Am. Nat., 183, 600–611.

Wood, S.N. (2011). Fast Stable Restricted Maximum Likelihood and Marginal Likelihood Estimation of Semiparametric Generalized Linear Models. J. R. Stat. Soc. Ser. B Stat. Methodol., 73, 3–36.

Wood, S.N. (2017). Generalized additive models: an introduction with R. chapman and hall/CRC.

