## Supplementary S1 for "Pollinator response diversity and turnover buffer pollination networks from habitat loss in human-modified landscapes"

### Table of contents:


- Table S1. List of pollinator species
- Table S2. List of plant species
- Table S3. Description of landscape variables tested.
- Table S4. Gams selection summary based on AIC
- Figure S1. Correlation per metric between different buffer scales
- Figure S2. Summary of response trends per pollinator guild

**Table S1.** List of pollinator species included in the study. We have analysed the data for species of the orders Diptera, Hymenoptera and Lepidoptera. Coleoptera species were also recorded but were removed from the analyses due to biological differences and lower expected movement.

| Pollinator guild | Family | Species | Nº of sites | Total records |
|---|---|---|---|---|
| Butterfly | Pieridae | *Pieris brassicae* | 22 | 82 |
| | Lycaenidae | *Lampides boeticus* | 7 | 26 |
| | | *Callophrys rubi* | 6 | 9 |
| | | *Lycaena phlaeas* | 5 | 16 |
| | Papilionidae | *Zerynthia rumina* | 5 | 13 |
| | Pieridae | *Colias crocea* | 4 | 10 |
| Hover fly | Syrphidae | *Episyrphus balteatus* | 20 | 103 |
| | Syrphidae | *Sphaerophoria scripta* | 14 | 31 |
| | Syrphidae | *Eristalis similis* | 8 | 12 |
| | Syrphidae | *Eupeodes corollae* | 6 | 7 |
| | Syrphidae | *Eristalis sp* | 5 | 6 |
| | Syrphidae | *Chrysotoxum intermedium* | 3 | 6 |
| | Platystomatidae | *Platynochaetus setosus* | 5 | 9 |
| Other fly | Bombyliidae | *Bombylius sp* | 11 | 47 |
| | | *Bombylella atra* | 6 | 19 |
| | | *Bombylius torquatus* | 4 | 5 |
| | | *Usia (Microusia) sp* | 3 | 19 |
| | | *Usia aenea* | 3 | 5 |
| | | *Dischistus senex* | 3 | 3 |
| | Calliphoridae | *Rhyncomyia cuprea* | 10 | 15 |
| | | *Rhyncomyia sp* | 6 | 12 |
| | | *Stomorhina sp* | 6 | 8 |
| | | *Calliphora sp* | 4 | 25 |
| | | *Lucilia sp* | 4 | 5 |
| | Muscidae | *Musca domestica* | 18 | 49 |
| | Syrphidae | *Merodon sp* | 11 | 15 |
| Wild bee | Andrenidae | *Panurgus calcaratus* | 20 | 72 |
| | | *Andrena sp* | 20 | 54 |
| | | *Andrena rhyssonota* | 8 | 20 |
| | | *Andrena morio* | 8 | 15 |
| | | *Andrena flavipes* | 8 | 14 |
| | | *Andrena alluaudi* | 7 | 18 |

| | | | | |
|---|---|---|---|---|
| | | *Panurgus sp* | 7 | 14 |
| | | *Panurgus banksianus* | 7 | 13 |
| | | *Flavipanurgus venustus* | 5 | 15 |
| | | *Andrena villipes* | 5 | 14 |
| | | *Panurgus cephalotes* | 4 | 14 |
| | | *Andrena granulosa* | 4 | 10 |
| | | *Andrena nigroaenea* | 3 | 12 |
| | | *Andrena abrupta* | 3 | 5 |
| | | *Andrena alma* | 3 | 5 |
| | | *Andrena colletiformis* | 3 | 4 |
| | Apidae | *Bombus terrestris* | 22 | 94 |
| | | *Xylocopa cantabrita* | 20 | 97 |
| | | *Anthophora bimaculata* | 13 | 36 |
| | | *Anthophora dispar* | 13 | 36 |
| | | *Eucera sp* | 11 | 21 |
| | | *Eucera collaris* | 9 | 46 |
| | | *Anthophora sp* | 8 | 16 |
| | | *Nomada sp* | 8 | 15 |
| | | *Anthophora plumipes* | 7 | 26 |
| | | *Anthophora retusa* | 7 | 17 |
| | | *Nomada merceti* | 7 | 12 |
| | | *Eucera clypeata* | 6 | 18 |
| | | *Eucera longicornis* | 5 | 9 |
| | | *Nomada succincta* | 3 | 6 |
| | | *Ceratina cucurbitina* | 3 | 5 |
| | | *Ceratina sp* | 3 | 4 |
| | | *Eucera nigrescens* | 3 | 4 |
| | | *Nomada agrestis* | 3 | 3 |
| | Colletidae | *Colletes acutus* | 6 | 25 |
| | | *Hylaeus pictipes* | 3 | 14 |
| | | *Colletes sp* | 3 | 3 |
| | | *Lasioglossum sp* | 21 | 70 |
| | | *Lasioglossum inmunitum* | 17 | 89 |
| | | *Lasioglossum prasinum* | 10 | 16 |
| | | *Lasioglossum albocinctum* | 5 | 10 |
| | | *Lasioglossum dusmeti* | 3 | 4 |
| | | *Sphecodes sp* | 3 | 4 |
| | | *Lasioglossum malachurum* | 3 | 3 |

| | | | | |
|---|---|---|---|---|
| | Megachilidae | *Megachile sicula* | 8 | 27 |
| | | *Osmia bicornis* | 4 | 5 |
| | | *Osmia sp* | 4 | 5 |
| | Melittidae | *Dasypoda cingulata* | 14 | 60 |
| | | *Dasypoda sp* | 6 | 18 |
| | | *Dasypoda crassicornis* | 6 | 9 |
| | | *Dasypoda morotei* | 6 | 9 |
| | | *Dasypoda iberica* | 4 | 6 |

**Table S2.** Plant species included in the study, the number of sites where each was recorded, and mean flower cover per occupied site (%).

| Plant species | Nº of sites | Mean flower cover per occupied site (%) |
|---|---|---|
| *Anchusa calcarea* | 2 | 1.00 |
| *Andryala integrifolia* | 9 | 11.61 |
| *Andryala sp* | 1 | 1.00 |
| *Anemone palmata* | 2 | 4.00 |
| *Asphodelus ramosus* | 2 | 0.92 |
| *Centaurea calcitrapa* | 1 | 5.00 |
| *Centaurea nigra* | 1 | 0.50 |
| *Cerinthe gymnandra* | 5 | 5.00 |
| *Cistus crispus* | 22 | 19.07 |
| *Cistus ladanifer* | 16 | 7.95 |
| *Cistus libanotis* | 10 | 10.42 |
| *Cistus monspeliensis* | 3 | 2.25 |
| *Cistus salviifolius* | 28 | 12.41 |
| *Crepis sp* | 12 | 5.21 |
| *Cytisus scoparius* | 4 | 4.88 |
| *Diplotaxis sp* | 2 | 3.00 |
| *Echium plantagineum* | 6 | 5.04 |
| *Echium sp* | 4 | 2.50 |
| *Echium vulgare* | 12 | 5.58 |
| *Erica umbellata* | 1 | 13.75 |
| *Erodium cicutarium* | 1 | 1.00 |
| *Erodium primulaceum* | 18 | 3.28 |
| *Galactites tomentosa* | 8 | 11.38 |
| *Genista hirsuta* | 7 | 7.43 |
| *Geranium molle* | 2 | 5.25 |
| *Gladiolus italicus* | 1 | 1.00 |
| *Glebionis coronaria* | 4 | 7.50 |
| *Halimium calycinum* | 22 | 5.66 |
| *Halimium halimifolium* | 22 | 13.03 |
| *Hypochaeris radicata* | 2 | 22.50 |
| *Jasione montana* | 2 | 2.00 |
| *Klasea nudicaulis* | 1 | 1.00 |
| *Lavandula pedunculata* | 17 | 9.72 |
| *Lavandula sp* | 4 | 11.38 |
| *Lavandula stoechas* | 24 | 8.11 |
| *Leontodon sp* | 6 | 4.17 |
| *Leontodon taraxacoides* | 13 | 12.08 |

| | | |
|---|---|---|
| *Linaria viscosa* | 14 | 3.80 |
| *Malcolmia lacera* | 15 | 4.32 |
| *Matricaria sp* | 5 | 5.20 |
| *Muscari comosum* | 3 | 2.17 |
| *Oxalis pes-caprae* | 2 | 6.50 |
| *Pallenis spinosa* | 1 | 8.50 |
| *Pistacia lentiscus* | 1 | 5.00 |
| *Raphanus raphanistrum* | 10 | 5.70 |
| *Reseda phyteuma* | 1 | 7.00 |
| *Ruta sp* | 1 | 0.50 |
| *Salvia rosmarinus* | 19 | 8.70 |
| *Silene colorata* | 2 | 3.50 |
| *Sonchus sp* | 5 | 2.10 |
| *Spartium junceum* | 2 | 11.25 |
| *Teucrium fruticans* | 5 | 11.27 |
| *Thapsia villosa* | 12 | 22.08 |
| *Tolpis barbata* | 10 | 9.15 |
| *Tuberaria guttata* | 9 | 15.50 |
| *Ulex australis* | 3 | 6.50 |

**Table S3.** Description of landscape variables tested. They involve three dimensions of the landscape: habitat quantity, habitat quality and habitat heterogeneity. In total we tested eight variables accounting for the patch characteristics and the type of habitat present. All variables are extracted for the 2km buffer size.

| Dimension | Variable | Description | Units |
|---|---|---|---|
| 1) Habitat quantity (natural habitat) | Percentage of natural habitat loss | Percentage of natural habitat within a 2 km radius around each site. | % |
| | Mean patch area | Mean area of habitat patches within a 2 km radius. | $m^2$ |
| | Largest patch area | Area of the largest habitat patch within a 2 km radius. | $m^2$ |
| 2) Habitat heterogeneity | Habitat Shannon index | Shannon diversity index of land-cover classes within a 2 km radius. | Unitless |
| 3) Fragmentation | Number of patches | Number of habitat patches within a 2 km radius. | Count |
| | Edge length | Total length of habitat edges within a 2 km radius. | m |
| | Edge density | Total edge length standardized by landscape area. | $m\ ha^{-1}$ |
| | Mean shape index | Mean shape complexity of habitat patches. | Unitless |

**Table S4.** Model selection results for alternative landscape metrics calculated within a 2 km radius around each sampling site. Each model included a general response shared across pollinator species, together with species-specific responses to the landscape metric. Models were fitted using a zero-inflated Poisson distribution and ranked according to AIC.

| Variable | AIC | ΔAIC | Akaike weight | Deviance explained |
|---|---|---|---|---|
| Percentage of natural habitat loss | 2239.65 | 0.00 | 1.0 | 0.51 |
| Edge density | 2269.21 | 29.56 | 0.0 | 0.49 |
| Largest patch area | 2286.72 | 47.07 | 0.0 | 0.47 |
| Habitat Shannon index | 2295.57 | 55.92 | 0.0 | 0.46 |
| Number of patches | 2296.75 | 57.10 | 0.0 | 0.45 |
| Mean patch area | 2326.21 | 86.56 | 0.0 | 0.42 |
| Edge length | 2355.53 | 115.88 | 0.0 | 0.40 |
| Mean shape index | 2387.19 | 147.54 | 0.0 | 0.34 |

**Figure S1.** Pearson correlations between landscape metrics calculated within 500 m and 1000 m buffers and their corresponding values calculated within a 2 km buffer around each sampling site. Correlations were generally moderate to strong, indicating that broad patterns of landscape composition and configuration were largely consistent across spatial extents.

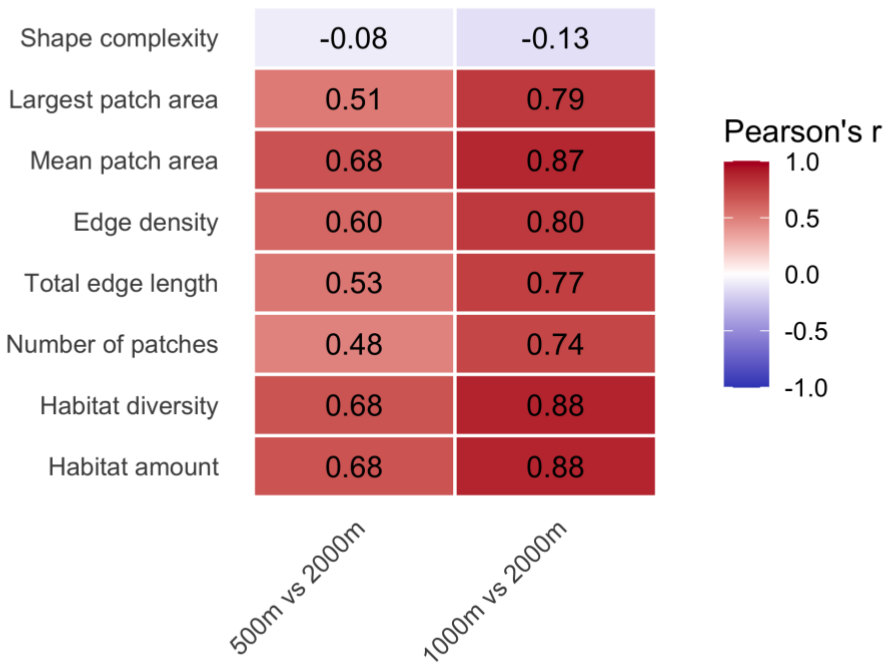

**Figure S2.** Predicted number of pollinator visits along the habitat loss gradient. Solid lines represent the mean predicted response of all taxa within each pollinator guild, while dashed lines represent individual taxon-specific trends. Overall, no pollinator guild showed a strong decline along the habitat-loss gradient, although hoverflies exhibited the most negative guild-level response. Wild bees showed the greatest diversity of species-specific responses, with a broader range of net trends compared with butterflies and other flies.

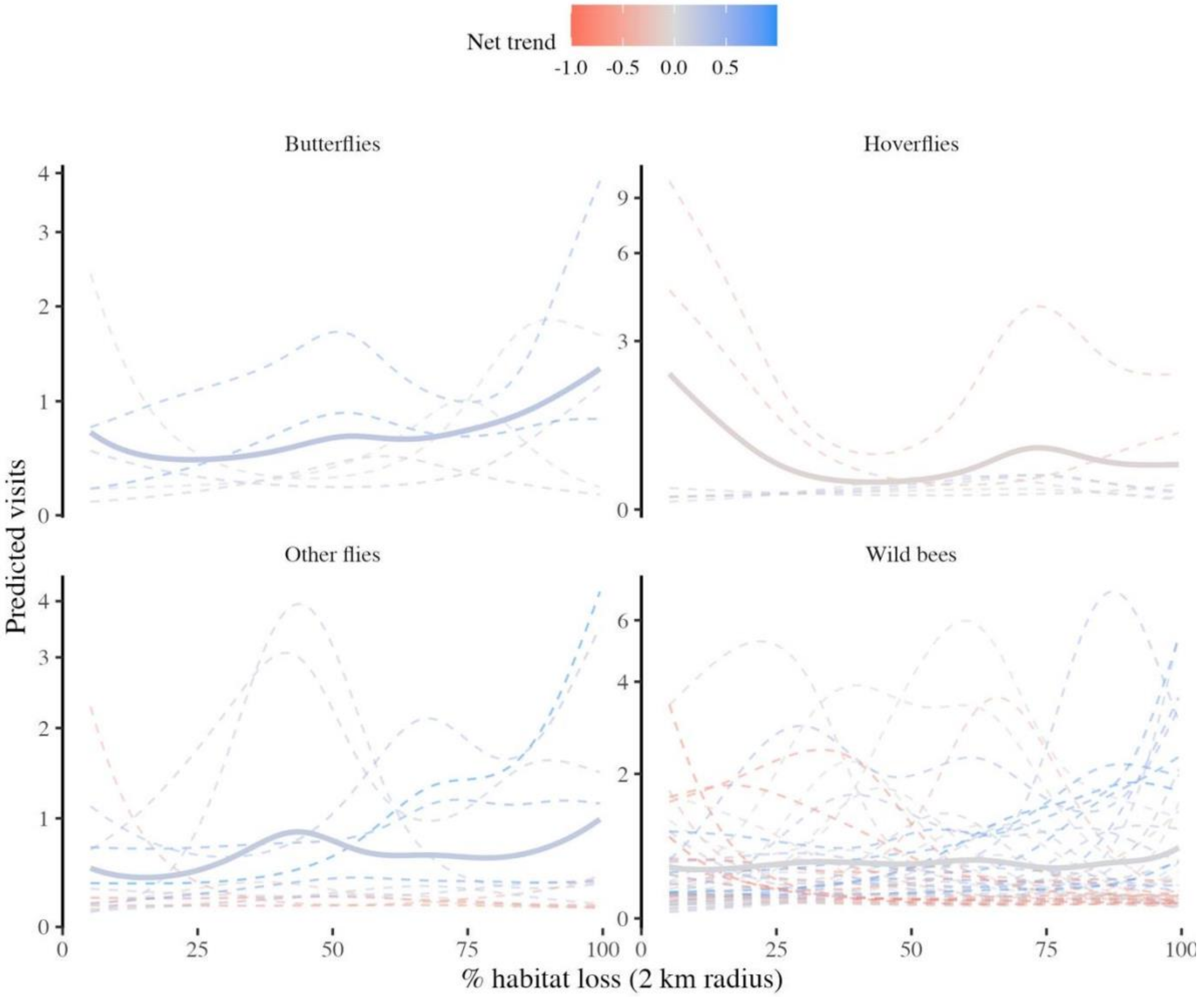
